# A computer vision approach for studying fossorial and cryptic crabs

**DOI:** 10.1101/2020.05.11.085803

**Authors:** César Herrera, Janine Sheaves, Ronald Baker, Marcus Sheaves

## Abstract

Despite the increasing need to catalogue and describe biodiversity and the ecosystem processes it underpins, these tasks remain inherently challenging. This is particularly true for species that are difficult to observe in their natural environment, such as fossorial and cryptic crabs that inhabit intertidal sediments. Traditional sampling techniques for intertidal crabs are often invasive, labour intensive and/or inconsistent. These factors can limit the amount and type of data that can be collected which in turn hinders our ability to obtain reliable ecological estimates and compare findings between studies. Computer vision and machine learning algorithms present an opportunity to innovate and improve sampling approaches. Moreover, cheaper and tougher recording devices and the diversity of open source software further boost the possibilities of achieving rigorous image-based sampling, which can broaden the range of questions that can be addressed from the data collected. Despite its significant potential, the software and algorithms associated with image-based sampling may be daunting to researchers without expertise in computer vision. Therefore, there is a need to develop protocols and data processing workflows to showcase the value of embracing new technologies. This paper presents a non-invasive computer vision and learning protocol for sampling fossorial and cryptic crabs in their natural environment. The image-based protocol is underpinned by fit-for-purpose and off-the-shelf software. We demonstrate this approach using fiddler crab and sediment recordings to study and quantify crab abundance, motion patterns, behaviour, intraspecific interactions, and estimate bioturbation rates. We discuss current limitations in this protocol and identify opportunities for improvement and additional data stream options that can be obtained from this approach. We conclude that this protocol can overcome some of the limitations associated with traditional techniques for sampling intertidal crabs, and could be applied to other taxa or ecosystems that present similar challenges. We believe this sampling and analytical framework represents an important step forward in understanding the ecology of species and their functional role within ecosystems.

## Introduction

Over the last century the rate at which species, habitats and ecosystems are being lost has become one of the most concerning challenges faced by humanity (Polidoro *et al.* 2010; Barnosky *et al.* 2011; Ripple *et al.* 2017). At no other time has it been more important to describe, quantify and understand patterns of species loss, and the relationship between species loss and the functioning of ecosystems (Cardinale 2012; Gamfeldt *et al.* 2015). A primary consequence of this urgency is the necessity to scale up the volume of relevant information gathered about species and ecosystems, and enable actionable and rapid scientific and policy response through rapid access to data stored in curated databases, for instance by using Technoecology (Allan *et al.* 2018) and Ecoinformatics (Michener & Jones 2012) approaches. For biologists and ecologists with dirt under their fingernails, this is an even more challenging task, as many traditional methods of sampling and collection are logistically limited by personnel-power (i.e. manual sampling, sorting, identifying, and measuring).

Working with cryptic and fossorial organisms, such as fiddler crabs, imposes additional challenges in achieving exhaustive biodiversity censuses and functional ecology assessments. The cryptic nature of some species makes it difficult to confirm species identities in the field. Furthermore, fiddler crab taxonomic classification and phylogeny remain a contentious topic (Crane 1975; Levinton, Sturmbauer & Christy 1996; Sturmbauer, Levinton & Christy 1996; Rosenberg 2001; Beinlich & Von Hagen 2006; Rosenberg 2014; Hsi-Te *et al.* 2016) which impedes species identification by ecologists and thwarts attempts to synthesise published literature to the level of species. Their fossorial nature coupled with their sensitivity to observers and fast speed also limits our ability to study these creatures which spend a considerable proportion of the time inside burrows (Caravello & Cameron 1991; Reaney 2007). Despite these limitations, fiddlers crabs as a group are among the most studied crustaceans (e.g. see references selection in Rosenberg 2001 introduction). Fiddler crabs, along with other intertidal and coastal crabs, regulate energy and matter fluxes in various ecosystems such as mangroves, salt marshes, sandy beaches and mudflats (Robertson & Daniel 1989; Sheaves & Molony 2000; Webb & Eyre 2004; Kristensen 2008; Nerot *et al.* 2009; Smith, Wilcox & Lessmann 2009). For instance, crab bioturbation plays a preponderant role in shaping soil composition, texture, and microbial community, which in turns drives nutrient and soil metabolism in coastal sediments (Gribsholt, Kostka & Kristensen 2003; Wang *et al.* 2010; Fanjul *et al.* 2011; Gittman & Keller 2013; Fanjul *et al.* 2015; Booth *et al.* 2019). The importance of crab bioturbation has been hypothesized to be relative to the total burrow volume and sediment turnover rate (Katz 1980; Gribsholt, Kostka & Kristensen 2003; Wang *et al.* 2010) which is proportional to crab density and burrow behaviour. Despite substantial evidence on the ecological function of fossorial intertidal crabs, caution is necessary when generalizing their importance because precise function is likely to vary depending on environmental conditions (Michaels & Zieman 2013; Natalio *et al.* 2017).

Despite the functional role of fiddler crabs, the methods used to estimate ecological measurements such as movement patterns, abundance, sex ratios, behaviour and bioturbation rate, among others, present some serious pitfalls and limitations (Nobbs & McGuinness 1999; Kent & McGuinness 2006; Schlacher *et al.* 2016). Invasive techniques, such as sediment excavation and digging (e.g. Colby & Fonseca 1984), installation of pitfall traps (e.g. Salmon & Hyatt 1983) and others, have the advantage of gathering the most detailed information about population structure (e.g. densities, sex ratios and size structure) and are assumed to offer the most reliable estimates (Lourenco, Paula & Henriques 2000; Macia, Quincardete & Paula 2001; Skov & Hartnoll 2001; Skov *et al.* 2002). However, these methods are time-consuming and modify the crabs’ natural habitat. Moreover, given the fossorial nature of fiddler crabs, some invasive techniques, such as pitfall traps, do not guarantee accurate and precise estimates across habitats or seasons because these are designed to monitor crab activity, instead of densities (Lee 1998). Alternative and widespread non-invasive methods, such as burrow counts (e.g. Mouton & Felder 1996; Lourenco, Paula & Henriques 2000) and distant counts of active individuals (e.g. Macia, Quincardete & Paula 2001; Skov & Hartnoll 2001; Jordao & Oliveira 2003), are only reliable for specific crab species, require site-specific calibration, and are based on assumptions about species behaviour, population structure and phenology (Macia, Quincardete & Paula 2001; Skov & Hartnoll 2001; Jordao & Oliveira 2003; Schlacher *et al.* 2016). These are likely to give a biased sample of the population (i.e. Hawthorne effect and other observation and sampling bias). Similarly, bioturbation rates estimates often require methods such as making burrow casts or quantifying the amount of sediment moved (e.g. Katz 1980; Escapa *et al.* 2007; Wang *et al.* 2010), which are hard to apply intensively and extensively. In contrast, while non-invasive methods, such as video recordings (e.g. Nordhaus, Diele & Wolff 2009), can minimize or overcome some of these challenges (i.e. biases), their application has been constrained because of the volume of data created and lengthy processing times. Thus, the fossorial lifestyle, behavioural habits and cryptic characteristics of fiddler crabs present an opportunity to innovate and to produce fit-for-purpose sampling techniques.

New technologies offer alternatives for streamlining the collection of biological and ecological data, while automation alleviates the bottlenecks in processing and analysis workflows. One of these technologies is machine vision, the use of hardware/software interfaces to develop computerized procedures and analyses based on images. Computer vision applications have penetrated the field of ecology and proven to be useful in extracting information from images, still or video (e.g. Zion 2012; Spampinato *et al.* 2015; Shafait *et al.* 2016; Villon *et al.* 2018; Weinstein 2018; Schneider *et al.* 2019). There are two main areas in which computer vision is accelerating image analysis for biologists and ecologists. Firstly, through the automation of the image classification and image segmentation, i.e., identification of features in images (i.e. organisms, habitats, organs, etc.) and the isolation of these features from other elements in the image (Harvey & Cappo 2001; Mathiassen *et al.* 2011; Matai *et al.* 2012; Li *et al.* 2015; Piechaud *et al.* 2019); and secondly, through automation of object tracking in image sequences, i.e., following the movement of features (Xie, Kham & Shah; Dell *et al.* 2014; Sridhar, Roche & Gingins 2019). These two computer vision tasks promise to reduce the amount of time spent processing image data, so easing the bottleneck created by collecting far more data than can be processed time-effectively. Unfortunately, computer vision solutions to biological and ecological problems are not always useful for more than one purpose, so, our field requires continuous development of heuristic and fit-for-purpose methods and algorithms.

Here we propose a workflow to collect biological and ecological metrics of intertidal crabs in mudflats and sandflats based on field video recordings. This workflow is underpinned by computer vision and machine learning algorithms which extract data on motion trajectories, colouration, size and bioturbation. A significant portion of this manuscript is devoted to describing fit-for-purpose methods, but we also list current challenges and future technological avenues to obtain additional biological and ecological metrics from image sequences. We believe that our approach can be useful for biologists and ecologists working in similar systems and taxa, regardless of the spatial and temporal scale of their work.

## Materials and Methods

### Study area, equipment and specimen collection

*Tubuca polita* (Crane, 1975) individuals were recorded in a mudflat of the Annandale wetland in Townsville, North Queensland, Australia. A sediment area with low vegetation coverage was selected to ensure maximum visibility of crabs during recordings. A virtual quadrat, 80 x 80 cm, was monitored for 30 minutes. Quadrat vertices were marked with circular coloured indicators inserted into the sediment. The coloured indicators were used as size and Cartesian coordinate references. To ensure that the methods were not capture device-dependent, several cameras were used during trials (e.g. GoPro HERO3, Garmin VIRB XE, Apple IPhone 8 and Canon 60D). All cameras recorded sufficient quality video to follow the workflow described below. Comparisons between cameras are not presented in this paper as it was beyond of the scope of this study. However, despite similar outcomes across different cameras, in a study situation, to maximize reproducibility, a single camera model should be used based on specific demands such as portability, price, image quality, and features (e.g. camera colour profile, lens distortion, focal length, frame resolution, frame rate, video compression and aspect ratio). To validate morphometric data and species identity from the videos, at the end of the recording period we collected as many of the observed crabs as possible. However, many crabs evaded capture and hence, individuals whose identity could not be confirmed were excluded from subsequent analysis.

### Image acquisition and analysis of crab recordings

To track and count individuals, and assess their size and colour, a GoPro HERO3 camera was positioned to fully cover the virtual quadrat area in the field of view. The camera position used was mode II (Fig 1-A) which minimizes disturbance to the natural vegetation surrounding the quadrat (i.e. mangroves with low canopy) and minimizes equipment shadows casting over the quadrat. Camera position mode I (Fig 1-A) is ideal for tracking individuals but normally involves stepping on (or close) to the area of interest which can increase crab burrow latency (personal observation; latency measured as the time of crab emergence from burrow after the observer abandons the area). Camera position mode III is ideal for assessing feeding rates and behaviour (Fig 1-A, e.g. see S1). Geographic position, sediment type, surrounding vegetation, observed tide, time of day, camera settings, and observed species composition were noted at the start of each video (Fig 1-B). To demonstrate the capacity of the analyses presented below these were only performed in the last five minutes section of the recorded video, when all crabs appeared to have resumed normal behaviour and after the maximum crab abundance was observed. Digital metadata and recordings were copied to a desktop computer, and three duplicates were created as backups (Fig 1-C). Several computer vision tasks were used during video analysis. Perspective correction was applied to guarantee the area per pixel ratio was similar across every region in the field of view (Fig 1-D-i). Individual crabs were isolated and movement was detected by using image segmentation (Fig 1-D-ii) and movement detection (Fig 1-D-iii) algorithms. Crabs were tracked in a composite video created by blending original frames with frames resulting from image segmentation (Fig 1-D-iv). Several challenges were found during the workflow described above (Table 1, Figure 1 D-vi, D-vii and D-viii). The most prevalent challenge was individual assignment when more than two crabs engaged in fights. When these challenges were encountered individual tracking was carried out manually.

**Table 1:** Challenges associated with identifying and tracking crabs in their environment: causes, effects and mitigation actions.

| Challenge | Cause | Effect | Mitigation action |
| --- | --- | --- | --- |
| Identify and track organisms in unconstrained environments. This is, environments with sudden and variable change in contrast and in illumination conditions. | Sunlight casting through canopy and understory foliage can produce hard contrast between fully illuminated and dark patches of sediment. This effect is exacerbated when moderate and strong winds moved the canopy and understory foliage, producing a frequent shift of these contrast shades in the sediment. | Increase inability to detect organismal motion and reduce successful identification and tracking.<br>It can bias scientist's decision about where and when to sample. | Sample in areas where this effect is absent, or sample at times when light intensity and illumination levels minimize this challenge.<br>Explicitly acknowledge sampling bias and assumptions.<br>Use more robust models to describe the background and foreground, such as cluster background models, texturized models, models based on neural networks, and others (e.g. Culibrk *et al.* 2007; Guo, Dai & Hoiem 2011; Zhu, Guo & Ma 2012; Kumar & Agarwal 2013). These models are often more computationally intensive.<br>Track organisms manually under unfavourable light conditions. |
| Maintain correspondence and consistency in organisms tracking ID when:<br>Organisms are occluded behind vegetation or other objects in the landscape (Fig 1-D-vi).<br>Two or more organisms were interacting adjacent to each other (Fig 1-D-vii).<br>Organisms moves rapidly (Fig 1-D-viii). | Camera, equipment configuration, and heterogeneous landscape prevents a direct view of the organism at any given time.<br>The correspondence between organisms and their trackers is normally established only using distance information between current and previous time.<br>Crabs exhibit burst motion which increase tracking ID mismatch. | Reduce the success of tracking one or multiple individuals. | Position camera and equipment to minimise blind spots in the field of view (e.g. mode I Fig 1-A).<br>Include additional information to resolve organism and tracker assignment. For instance, colour, size, texture, speed and direction, among other variables, can be explicitly added and considered in the tracking model and assignment algorithm.<br>Avoid skipping or dropping frames during analysis. |

**Figure 1:**
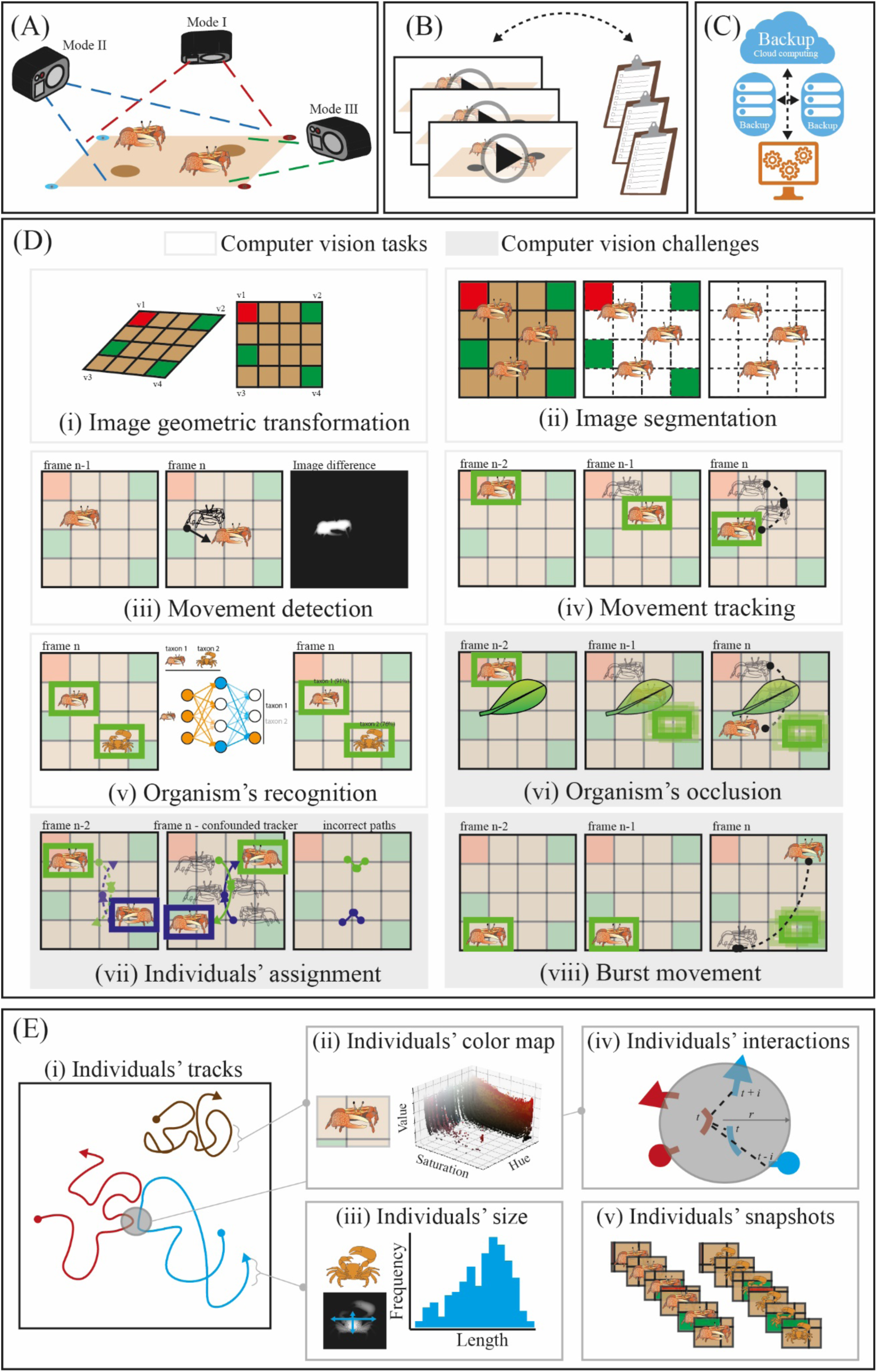
Schematic workflow for tracking, counting and measuring individual size and colour from videos. (A) Images can be captured from different viewpoints depending on the purpose of the study: mode I (vertical) and II (∼45°) are ideal for tracking, while mode III (horizontal/ground level) is convenient for observing behavioural displays. (B) Raw data (images) and metadata are linked and cross-referenced, and (C) data is backed up to multiple local and remote locations. (D) Common computer vision tasks applied and challenges encountered in image processing. (D-i) Image perspective transformation is useful to correct lens distortion and image composition. In natural settings foreground and background segmentation (D-ii) and movement detection (D-iii) are useful algorithms to isolate key features or organisms, and detect motion in images. Once features, objects or organisms are identified, tracking (D-iv) is done by estimating or predicting motion vectors among successive images. Furthermore, organisms’ classifier and pattern recognition (D-v, e.g. supervised, unsupervised, etc.) can aid and improve the performance of computer vision tasks. (D-vi) Organisms’ occlusion and features overlap reduce tracking success depending on the obstruction area and time. When multiple organisms interact in close proximity, individuals’ assignment during tracking is reduced potentially producing wrong track paths. That is, individuals’ identities were not preserved along the tracking period. (D-viii) Sudden and rapid motion can potentially produce loss of tracking. (E) Workflow results and products include individuals’ movement paths in the field of view (E-i). Knowing an individual’s position in the field of view at any time allows us to extract additional information such as individual colour and colour change (E-ii), size (E-iii) and number of close interactions with other individuals at any time (*t*) for a given time window (*i*) and minimum distance (*r*) (E-iv). In addition, organisms’ snapshots (E-v) can be saved for training image classifiers.

The above computational methods are enabled in Crabspy (Herrera 2020), a fit-for-purpose programming tool developed to sample crabs. Crabspy is a heuristic, open source toolbox created using Python programming language (Python Software Foundation) and the libraries OpenCV (OpenCV 2015), Pandas (McKinney & others 2010), Scikit-Image (Van der Walt *et al.* 2014), Numpy (Van der Walt, Colbert & Varoquaux 2011) and Scikit-Learn (Pedregosa *et al.* 2011). Crabspy can be accessed and downloaded from its official repository in GitHub (https://github.com/CexyNature/Crabspy). Crabspy combines common image segmentation and machine learning algorithms to track crabs in videos. Essentially, in its current version, a tracker (i.e. bounding box) is seeded at the original position of the target individual, then the tracker position is updated based on the movement of the individual in successive frames. The bounding box update can be done using any of the tracking methods available in the library OpenCV (i.e. MIL, BOOST, and KCF) or tracking can be done manually by the user. Crabspy detects movements and segments moving objects from the image to increase track reliability. The minimum relative size of objects moving, i.e. crabs, can be set using eroding and dilation parameters (i.e. kernel size). At any given frame, the track bounding box enclosing the target individual obtains its relative position, colour and size. Crabspy also has the flexibility to save images from individual crabs along successive frames.

During analysis, each individual was manually assigned an ID name, and its sex, handedness and species was recorded with the respective positional data (Fig 1-E-i). This step was performed manually, but with the increasing image collection two automated models able to assign sex and handedness were developed (i.e. model using a Support Vector Machine on Histogram of Oriented Gradients, and Convolutional Neural Networks, see S2). Crabs colouration can be extracted for any individual and any given time during the recording (Fig 1-E-ii). Individual colour data is useful to study species that exhibit whitening or brightening, or other changes of colouration (e.g. Hemmi *et al.* 2006; Takeshita 2019), in response to environmental and behavioural factors. The size of crabs at each frame was estimated by measuring the size of the Binary Large Objects (BLOBs, i.e. crabs shade) created after background removal and motion detection algorithms were applied. Each BLOB at each frame was measured along its two longer axes, which aimed to represent the width and length of the crab. These measurements resulted in a distribution of sizes per individual, from which central point estimates or confidence intervals could be estimated (Fig 1-E-iii). For each individual crab, a file containing position and size per frame, individual ID, species, sex and handedness was created. All files were combined and analysed using R software version 3.5.0 (R Core Team 2018) using libraries adehabitat (Calenge 2006), hab (Basille 2015), igraph (Csardi & Nepusz 2006), ggraph (Lin Pedersen 2018), and tidyverse suite (Wickham 2017). Crab densities, sizes and tracks were transformed from pixel units to centimetres by scaling based on the known side dimensions of the quadrat. Using the positional data from all individuals through time we extracted events where proximity between pairs of individuals was less than 10 cm within a two second time window. The proximity distance and time window can be adjusted depending on the question and working hypothesis (Fig 1-E-iv). For *T. polita*, we observed that a proximity of 10 cm or less is a good indicator of likely interactions between individuals. The type of interaction, agonistic and non-agonistic, was also recorded by observing each interacting pair in the video. Thus, a network analysis based on proximity was conducted to evaluate crab interaction patterns. In the network analysis, nodes represent individual crabs, and edges, or connections among nodes, characterize the type and degree of interaction among any pair of individuals. Finally, snapshots per individual at each frame were generated (Fig 1-E-v). These images were used to create training sets for image classification (Fig 1-D-v). Preliminary models, not presented in this paper, have achieved an 86% identification success rate for unseen crabs from the same population of genus *Tubuca* by transfer learning using the Single Shot Detector architecture (Liu *et al.* 2016) and the MobileNet Tensorflow model trained on the COCO dataset (Lin *et al.* 2014). As more crab pictures from this and other genera are gathered we believe successful identification rates are likely to improve. We suggest the utilization of deep neural network architecture trained on datasets with features that improve transfer learning for organismal identification (e.g. Van Horn *et al.* 2018).

### Image acquisition and analysis for 3-dimensional reconstructions of sediment

We explored the bioturbation activity of *T. polita* individuals by assessing the amount of sediment moved over time. Sediment moved was measured as the volume difference between two times. Volume difference was computed from 3D sediment reconstructions created using Visual Structure from Motion a photogrammetry technique. A permanent quadrat, 50 by 50 cm delimited with Ground Control Points (GCPs) in its vertices (Fig 2-A), was placed in the upper intertidal area of the muddy bank. This area also satisfied the following requirements: (1) presence of *T. polita* was confirmed, (2) no other crab species were present, and (3) the area was not flooded during and between samplings. Moreover, the sediment was sampled in low tide so there was no sediment transportation due to tidal change. The quadrat was independently recorded twice at two time points, namely time before and time after (7 days later), for a total of four sediment recordings. Each pair of independent video recordings at each time were filmed within minutes from each other. Thus, we assumed no change in sediment for these pairs which allowed us to calculate precision. The sediment was recorded using a camera Garmin VIRB XE by progressively moving the camera from side to side at constant speed (Fig 2-A). Videos were backed-up prior to analysis to reduce the likelihood of data entropy (Fig 2-C). Video scans normally produced videos no longer than 80 seconds. Image blur was minimized by setting the camera to a high frame rate and scanning the surface at times of maximum light. Meta information associated with each scan were recorded, and file management was performed as described above. Every 6^th^ frame was extracted from the videos resulting in 200 to 600 images for each video scan. These images were used to create photogrammetry reconstructions using the open-source and free software VisualSFM (Wu 2011; Wu 2013). VisualSFM and its library bindings find, match and triangulate features across images, and compute image cluster for creating sparse and dense reconstructions (Furukawa *et al.* 2010) (Fig 2-E-i). Dense cloud data was imported in the open-source and free software CloudCompare v2.20.2 Zephyrus (CloudCompare 2019). Mesh reconstruction, as well as textured, coloured and ortho-rectified 3D models can be created in CloudCompare (Fig 2E-ii, 2-E-iii). Our workflow calculations were done on dense point cloud data which preserve the inherent density of the sediment models (Fig 2-E-i). Point cloud datasets were ortho-rectified by scaling, rotating and transforming 3D clouds based on the known dimension and distance of GCPs, so the scale dimension of digital models coincided with the centimetre scale from the quadrat. Our analysis focused predominantly on computing volume differences from sediment scans, but as quality control and pre-analysis step we also evaluated that all scans presented similar surface density and roughness. Other topography metrics can be calculated as linear combinations of the eigenvectors and eigenvalues obtained from the 3D tensors for each point and for a given neighbourhood size (Weinmann, Jutzi & Mallet 2014; Hackel, Wegner & Schindler 2016). Digital elevation models were generated and cloud to cloud distances among all cloud pair combinations were calculated using the nearest neighbour distance and the quadratic function methods for accounting for density differences between point clouds. These two methods produced similar results (CloudCompare 2019).

**Figure 2:**
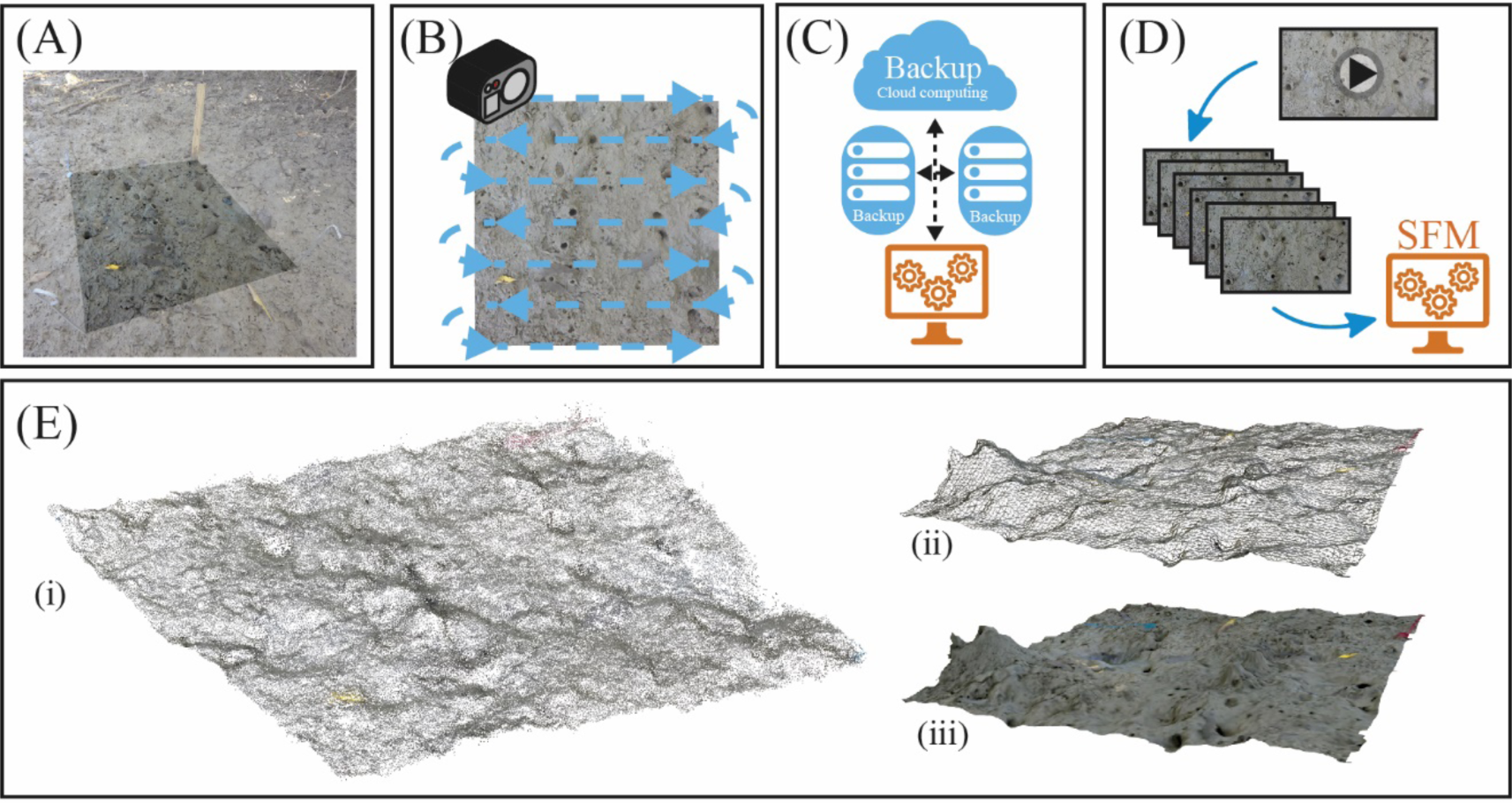
Schematic workflow for assessing sediment changes. (A) An area of 50 cm by 50 cm was defined in the sediment. Vertices were marked with coloured Ground Control Points (GCPs). (B) The area was scanned using a camera while recording a video. (C) Multiple copies from videos were created and saved in local and remote independent storage to prevent data entropy. (D) Frames from video were extracted and saved, and these were used to create photogrammetry reconstructions through Structure from Motion (SFM). From photogrammetry reconstructions three main data products can be created: dense point cloud data (E-i), mesh data (E-ii), and textured and coloured 3D models (E-iii). All these data types were ortho-rectified by using the GCPs.

## Results

*T. polita* individuals emerged from their burrows a few minutes after the sediment was exposed to air following the receding tide. From the 25 individuals observed in the field inside the quadrat, 17 were tracked during video analysis, and from these only ten were subsequently captured after the video recording was completed. Crab motion ranges and utilization areas overlapped among several individuals (Fig 3). Fourteen of the 17 tracked individuals were confirmed as males while for the remaining 3, sex was not determined. Ten individuals were right handed and three left handed. In four individuals, handedness could not be assessed because it was not verified in the field nor on the video. Total distance travelled per individual was variable (114.8 ± 69.16 cm), with a minimum of 24 cm (Fig 3, crab_6) and maximum of 240 cm (Fig 3, crab_3). In general, there were two types of individuals: those who wandered and those who stayed close to a burrow or territory. Some wandering individuals left the field of view during filming. *In situ* observations confirmed that at least two males, which left the field of view of the camera, moved up to four metres away from their original positions. Wandering crabs who stayed in the camera field of view travelled up to 161 cm (Fig 3, crab_10). Thus, some crabs attached to burrows accumulate longer travel distances that wandering crabs with higher distance displacement. For most non-wandering individuals the position of the home burrow defined the gravitational centre of the individual’s utilization area. The total distance travelled was higher in crabs with adjacent neighbours. Among this subset of individuals, short distance explorations to a neighbour’s burrow were observed. Burrow utilization was observed for eight individuals, and they actively defended their burrow from intruders.

**Figure 3:**
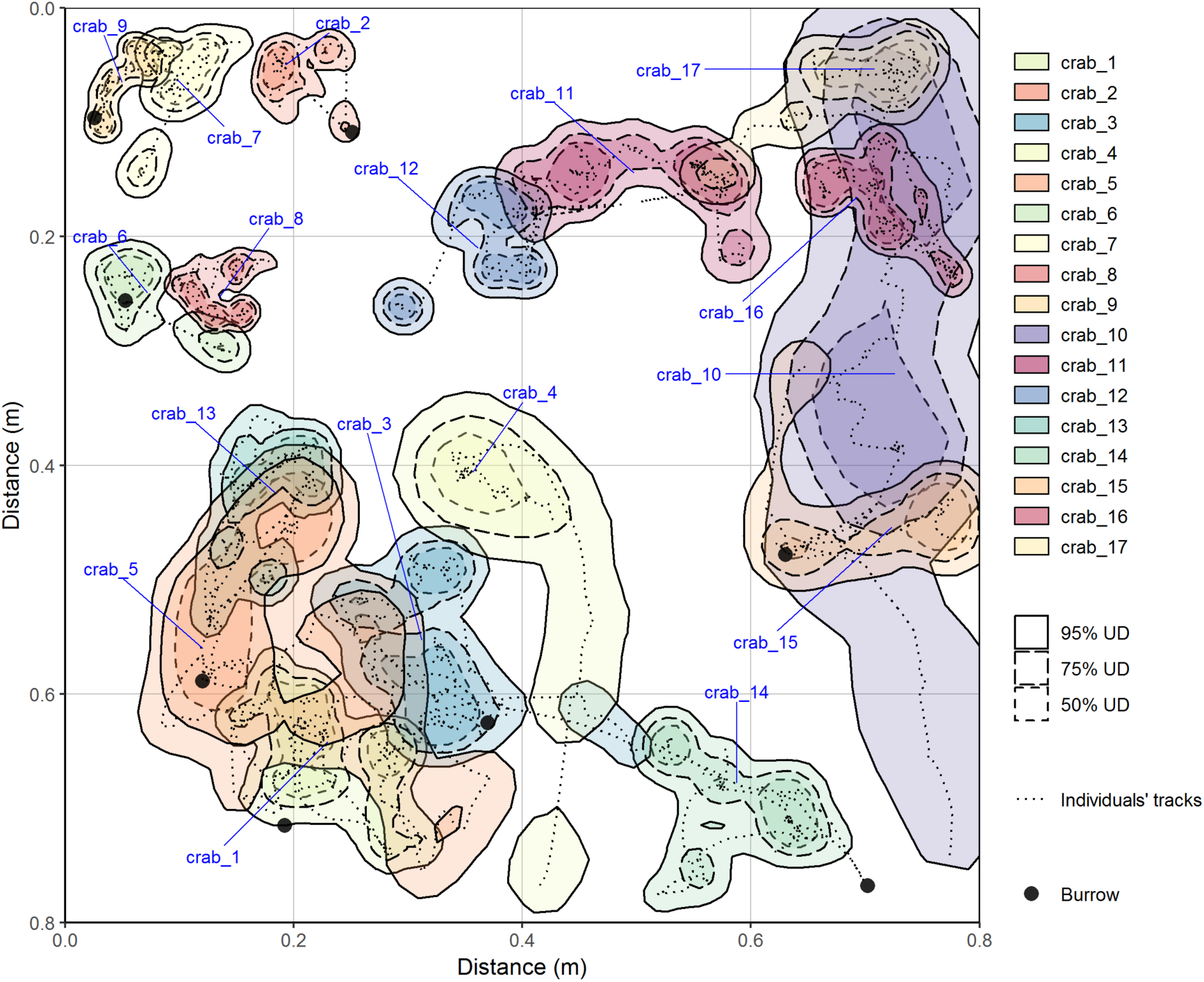
Movement tracks and kernel utilization distributions (50, 75 and 95%) for seventeen Ocypodidae crabs (*Tubuca polita*) observed during five minutes on an intertidal mudflat (80 cm by 80 cm area). Black circles represent burrows used during the observation period. Notice not all crabs remained within the quadrat for the entire observation time.

The size estimate distribution per individual produced a bimodal distribution in most individuals (Fig 4-A). This was expected as the longest two axes from each blob per frame were recorded. Various factors affected the size estimation. Firstly, the size of the kernel used for eroding and dilation during analysis constrained the possible values both axes could take. Erosion and dilation are two common computer vision filters to reduce noise. Thus, the axes’ lengths are a function of the kernel sizes. This factor affects the accuracy of the estimate. Secondly, our size estimate was also affected by waving, fighting, pushing or any other behaviour which changes the apparent size of the crab. Thirdly, individual close interaction to any other moving element in the landscape such as other crabs or vegetation overestimated size. Finally, long stops (lack of motion) underestimated sizes, as the blob progressively fades away (i.e. reduces its size) when movement is not detected. These last three factors affected the precision of the size estimate. The frequency of extreme deviations in the estimate due to the above factors is proportionally low given that a crab size estimate is generated for each frame in the video (Fig 4-A). We explored how well this size estimate predicted the carapace width and propodus length in the ten individuals that were captured and measured in the field (Fig 4-B). Even though some crabs frequency distributions for the size estimate presented two local maxima, we used the mean of the crab size estimate distribution as the predictor and the measured carapace width and propodus length as response variables and surrogates of crab size. While the first sampling moments for each mode (i.e. mean for each peak) would ideally represent crab carapace width and carapace length and therefore would be better estimators, we found that the first sampling moment (i.e. overall mean) for each distribution better represented the crab size after taking into consideration the above factors affecting the estimate. Assumptions for a linear model between size estimate and propodus length were not met. The linear model predicting carapace width based on our size estimate explained 65% of variation and this method tended to overestimate crab size (Fig 4-B).

**Figure 4:**
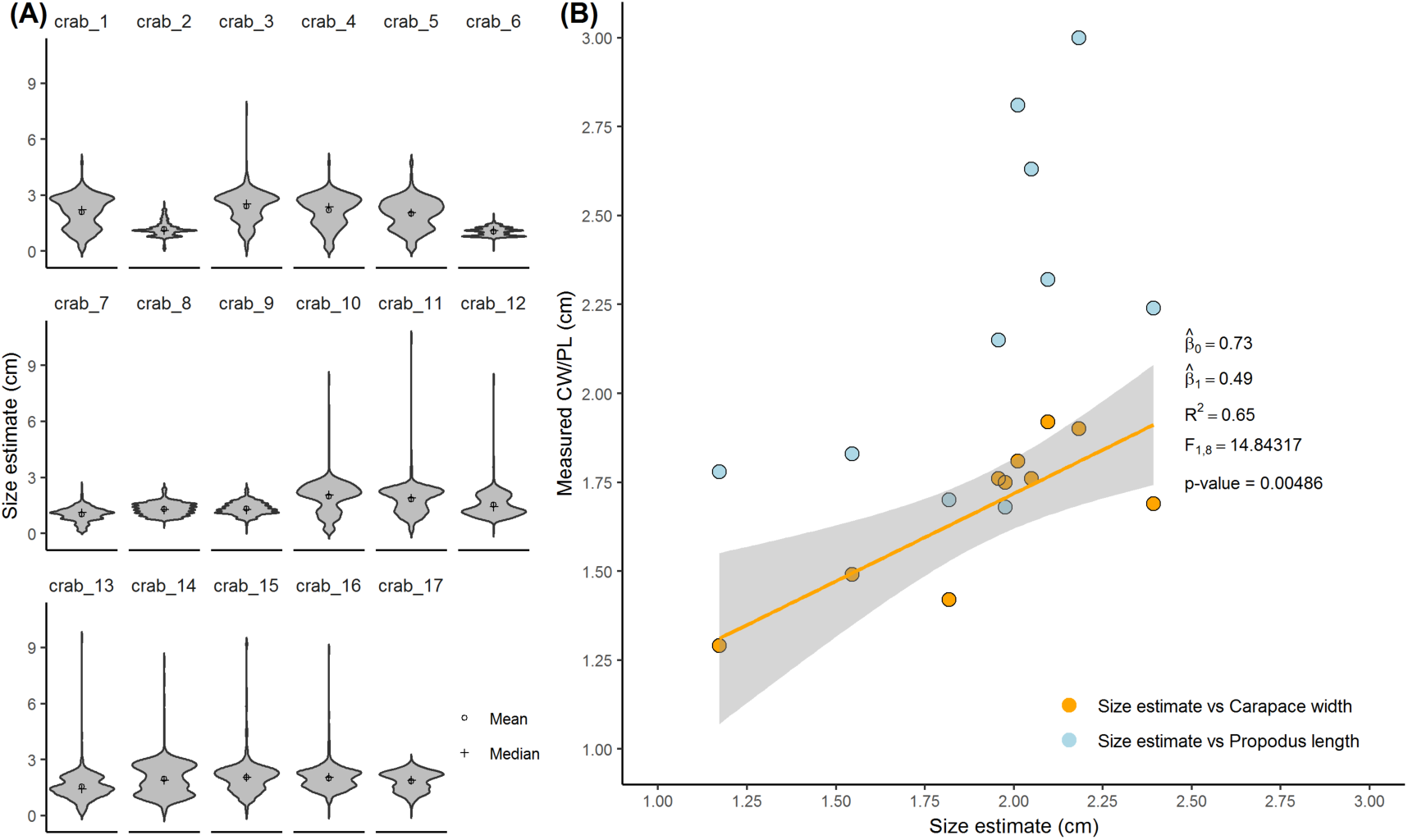
Size estimates, and size estimates and morphometric measurements relationship for *Tubuca polita* individuals. (A) Distribution of size estimates for the seventeen *Tubuca polita* individuals. Distribution is shown as violin plots to better visualize the histogram distribution for these seventeen individuals. Mean and median of these distribution are presented as circle and cross respectively, and these were assumed to represent the individuals’ carapace width point estimates. These distributions are continuous, but size estimate values are constrained by any limitation imposed in the blob algorithm. Thus, erosion and dilation kernel sizes affect the size estimates. Size estimations can greatly deviate from individuals’ carapace width in certain circumstances such as when crabs waved their claws, and while interacting close to other crabs or to any other moving element in the background (i.e. any other blob). (B) Size estimate (mean) versus morphometric measurements, i.e. Carapace Width (CW) and Propodus Length (PL), for ten out of the seventeen individuals. The fitted linear model (orange continuous line) describes the relationship between the size estimate and the measured CW (95% confidence interval is shown as gray shaded area).

The number of interactions between crabs varied on an individual basis (Fig 5). Most interactions occurred in the vicinity to a burrow, and most interactions involved at least one resident crab and one or more intruders or neighbours. Crab number 3 exhibited the highest number of interactions and the most number of agonistic interactions with different individuals (Fig 5). Some crab pairs kept a distance of 10 cm or less for a large proportion of the observed time, i.e. pairs 9-7, 6-8 and 12-11 (Fig 5). The number of edges in the network (19) is well below the possible number of edges for a proximity network with 17 individuals (i.e. 136). Size and handedness seemed to have an effect on the network structure, but this must be explored with larger data sets. As it has been observed in other systems (Barabasi & Albert 1999), some individuals acted as hubs, concentrating a higher proportion of edges (heterogeneous, scale free networks). The three network communities differed in size, with 8, 6 and 3 nodes.

**Figure 5:**
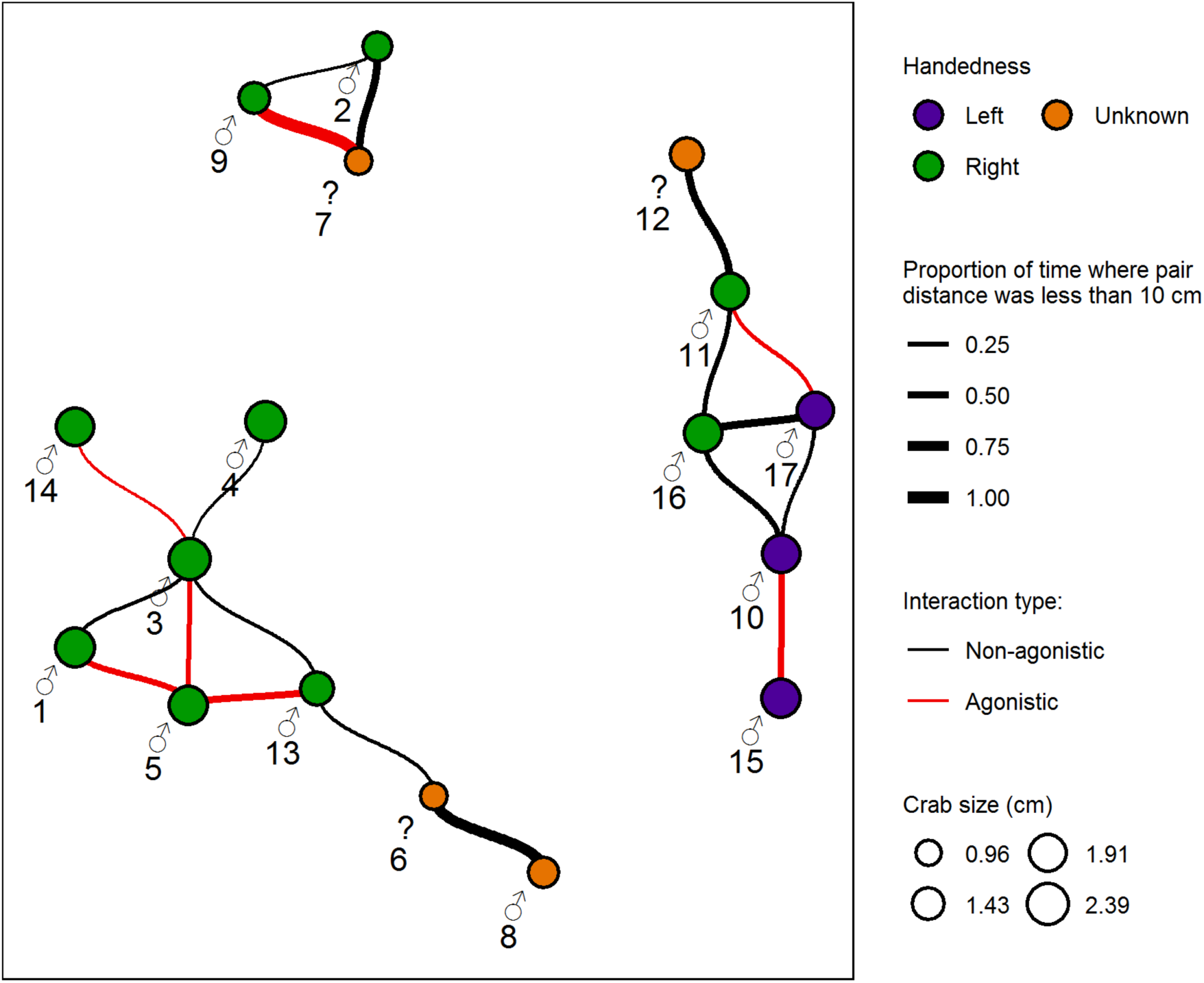
Interaction network for *Tubuca polita* individuals (nodes or n = 17; edges or *k* = 19). This undirected and weighted network was constructed from proximity data among pairs of individuals observed during five minutes. The node labels represent individuals ID and sex. Question marks indicate individuals for which sex was not confirmed. Node colour and size represent handedness and size estimates. Edges connecting nodes indicate that the distance between these pair of individuals was less than 10 cm in a two second window at least once in any period during the observed time. Edge width shows the proportion of time these individuals were within 10 cm from each other. Edge colour shows the type of interaction observed: red at least one agonistic, or fighting, or claws interlocked or pushing interaction; black at least one non-agonistic, or warning behaviour, or orientation change, or change of direction, or change on motion state, or no observable response to other individual (i.e. just proximity less than 10 cm).

The calculated volume difference between before and after was 1362 cm^3^ ± 47 (SE) over 0.25 square metres in a seven-day period. The volume difference between time replicates were 280 cm^3^ (before-before) and 171 cm^3^ (after-after). This represented a 13-20% precision error. Precision might be affected by small differences in point clouds which can be caused due to subtle differences in the image quality during capture and/or during the image analysis (Dandois, Olano & Ellis 2015; Bryson *et al.* 2017; Forsmoo *et al.* 2019). In fact, small differences were observed among the pair of digital elevation models taken a same time points (Fig 6-A, top row time before, bottom row time after). The distance between each point in a model and its homologue in the second model was expected to be zero when comparing models within the same time. Time replicates produced distance values predominantly equal to zero or close to zero (Fig 6-B), with less than 5% and 10% of cloud to cloud distances being greater than 0.4 centimetre for before-before and after-after comparisons, respectively. The four possible comparisons among different time models produced similar results, and these represented sediment change between time before and time after.

**Figure 6:**
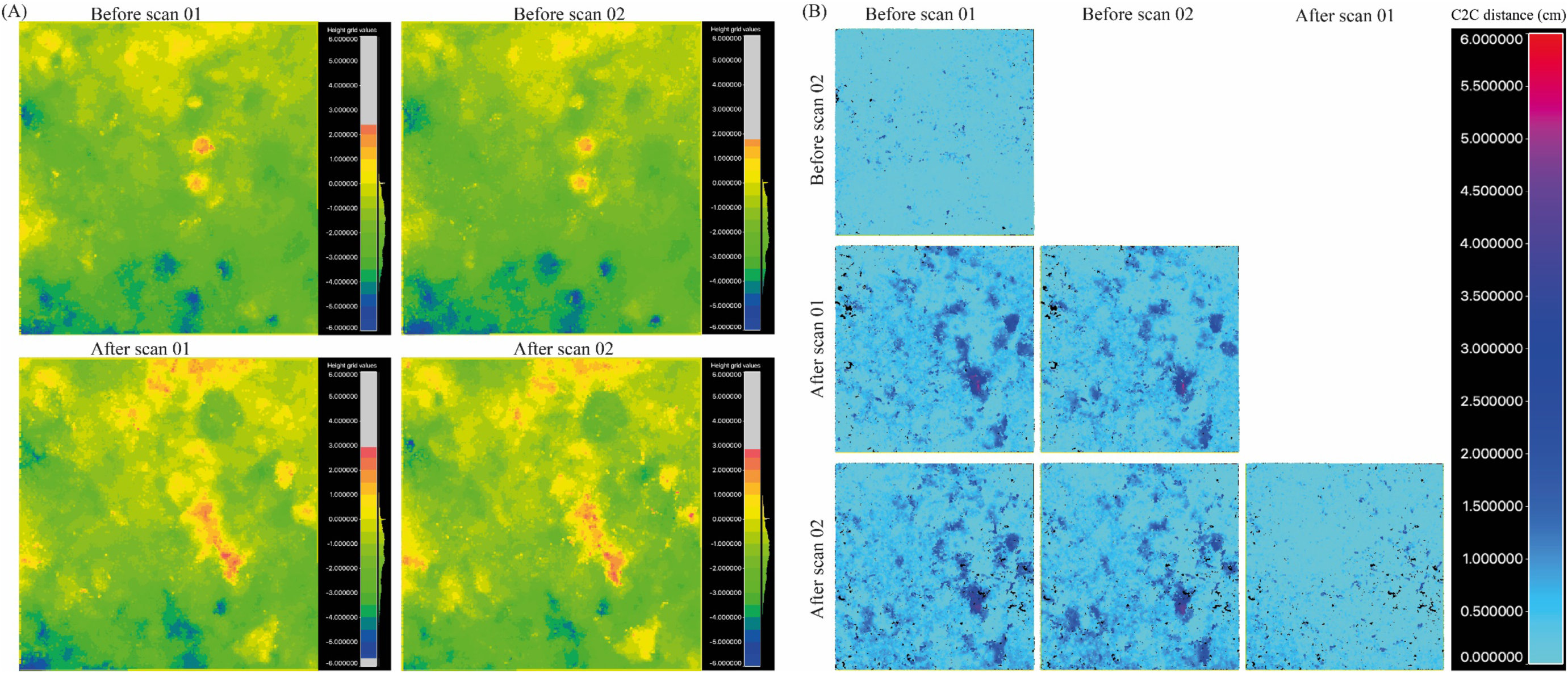
Comparison of raster generated from point cloud data for multiple sediment scans representing two times (before and after). Each time was independently scanned twice. (A) Digital elevation model in centimetres. Subtle differences can be observed within time (rows), these differences can be attributed to the number of features and cloud density for each data set. (B) Raster of computed cloud to cloud (C2C) distances among four scans. Values represent the calculated distance in centimetres among clouds using a quadratic function. Within time comparisons yield values close to zero, while among time comparison shows change in sediment.

## Discussion

We believe this is the first time that a non-invasive, cost- and time-efficient data workflow underpinned by computer vision has been used to gather rich data on benthic and fossorial crabs with potential scalability (Table 2). Our approach, using Open source and fit–for-purpose heuristic software, facilitates the rapid collection and processing of ecologically valuable data on small invertebrate organisms. For instance, here we focused on intertidal fiddler crabs, but our approach could easily be extended to include other taxa with similar modes of life. Although some of the techniques have been used before, to our knowledge they have not previously been used in combination to allow the collection of a wide range of information on an invertebrate species. Access to rich and big data overcomes issues associated with traditional sampling techniques such as hard to maintain methodological assumptions (Schlacher *et al.* 2016) and time- and labour-intensive costs. Moreover, such technological advances increase the amount of information of the species studied, thus these can accelerate our understanding of the ecology and biology of small species, which in turn can improve species and ecosystem management strategies.

**Table 2:** Type of data obtained using the Crabspy workflow, inventory of potential uses and opportunities for improvement.

| Data type | Domain knowledge | Application | Opportunity for improvement |
| --- | --- | --- | --- |
| Position, image and activity state | Animal motion | Characterize species movement. | Overcome tracking challenges described in table 1.<br>Incorporate and synchronize environmental sensors (e.g. temperature, humidity, light intensity, and other sensors) with video data. |
|  |  | Study individual self-orientation mechanisms in space and time. |  |
|  |  | Study utilization of space. |  |
|  | Community and population ecology | Assess species abundance. |  |
|  |  | Assess population demographics: size structure, sex ratio and handedness ratio. |  |
|  |  | Characterize intra and interspecific interactions. |  |
|  | Functional ecology | Assess bioturbation and feeding rates. |  |
|  |  | Understand metabolic and stoichiometry processes. |  |
|  | Ecophysiology | Study thermoregulation and water loss control patterns. |  |
|  | Behavioural ecology | Characterize waving, courtship and aggression displays through time. |  |
|  |  | Evaluate foraging behaviour and feeding preferences. |  |
|  |  | Study of communication and signalling |  |
|  |  | Understand of phenology and activity budgets. |  |
| Colour | Behavioural ecology | Study colour signalling. | Improve light measurement by using multispectral cameras or imaging spectrometer. |
|  |  | Study crypsis. |  |
|  | Evolution | Evaluate phenotype patterns and phenotype selection. |  |
| Sound and vibration | Ecophysiology | Study energy demands and trade-offs of stridulation at the individual level. | Incorporate sound wave recorder, phonograph, geophone or vibrometer to camera setup; and synchronize sound/vibration measurements with video. |
|  | Behavioural ecology | Study of stridulation mechanisms, patterns and functions at population and community level. |  |
|  |  | Study of acoustic communication |  |

Our 5-minute trial video analysis combined with 3D models demonstrates the considerable potential of employing computer vision techniques. This method allows us to obtain movement patterns of several individuals of *T. polita*, estimate their size and characterize their intra-specific interactions. Few studies have tried to characterize the movement patterns of fiddler crabs (Salmon 1984; Salmon 1987; Viscido & Wethey 2002). To our knowledge this is the first time that individual movement paths of multiple individuals in a natural setting has been obtained for any fiddler crab species. In this case, movement path information is complemented with information about the size, sex and handedness of the focal individual and neighbours. In addition, by categorizing behavioural displays throughout the video, sequential changes in behaviour can be related to specific motion and navigation changes. The need for such complementarity data, stressed by Nathan *et al.* (2008), makes this method and study system a good candidate to gather data to test mechanistic models of animal movement.

We have also estimated the bioturbation rate as volume change per area and time unit (778.29 cm^3^ per m^2^d^-1^). Further extensive sampling is required to assess the spatial and temporal patterns of *T. polita* bioturbation. Reported bioturbation rates for another fiddler crab species, *Minuca pugnax* (324 cm^3^ per m^2^d^-1^) are lower than our estimates (Katz (1980). However, other studies have reported greater volume change in other crab burrowing species such as *Helice tridens* (11484 cm^3^ per m^2^d^-1^)(Takeda and Kurihara (1987). Our measurement assumes that no other factor than fiddler crabs contributed to volume changes, and that volume changes are a suitable surrogate to sediment turnover by crabs (i.e. mass, Table 3). Although crab bioturbation is inherently related to crab density, burrow volume, species size and species burrow behaviour, the method we propose can be calculated independently to these other variables. Regardless of the method used all these assumptions (Table 3) must be carefully evaluated depending on the area and time of sampling. We believe the method proposed here to estimate bioturbation is reliable, reduces the potential negative impacts of destructive sampling in the habitat on crab populations, precision can be consistently calculated and known, and relies on similar assumptions as the methods employed till now. The proportionally high error in this method (13-20%) can potentially be reduced by artificially illuminating the areas scanned, and improving orthorectification and alignment procedure during analysis. Nonetheless, a full study of the error sources in this method and application will be worthy (see e.g. in Bryson *et al.* 2017).

**Table 3:**
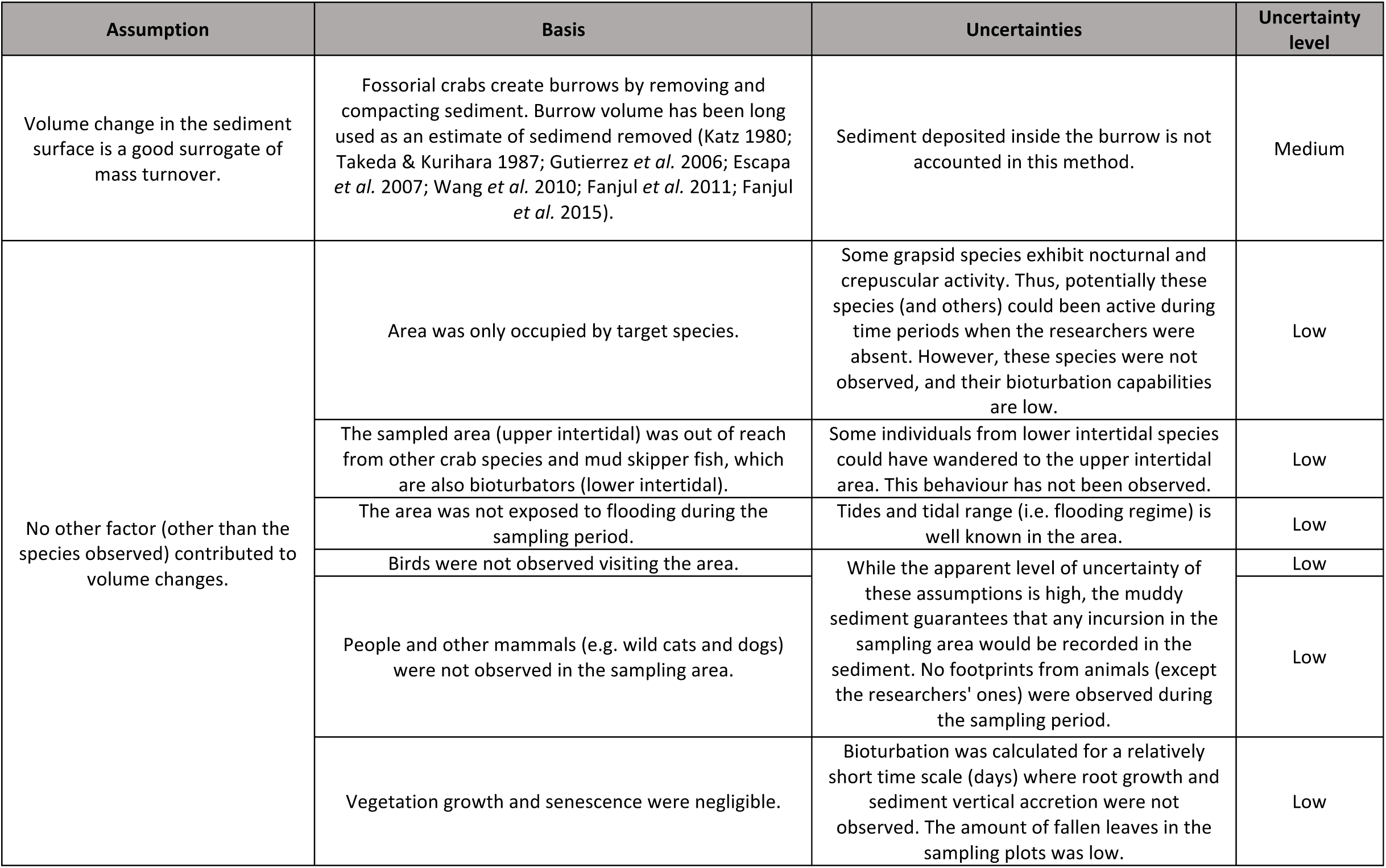
Assumptions underpinning bioturbation rates estimates.

There is a wealth of future opportunities that could be incorporated to extend the amount and type of data collected and analysed (Table 2). As more videos and images from target species are collected computer vision and machine learning models will likely become more robust and precise in identifying and predicting species behaviours. For instance: 1) automatically assessing frequency, time and pattern of waving and feeding activities; 2) automatically assigning sex, handedness and taxa using an image classifier; 3) extracting or generating additional spatial layers for the observed areas such as vegetation and sediment types; and 4) incorporating an elevation model in the tracking data, and incorporating and analysing information from inexpensive sensors such temperature and audio. Crab images captured during the current workflow together with the fine-tuning of algorithms already available will make it possible to achieve 1 to 4. For instance, number 1 and 2 have been already done using training obtained following workflow from this paper (S2). Number three requires more sophisticated segmentation techniques able to isolate vegetation and sediment layers, and/or the use of multi-spectral cameras which can facilitate the acquisition and isolation of these layers. Number four requires simple integration of tracking data to the elevation models which is mainly a computational tasks. Furthermore, keeping original videos or images, will allow us to retrospectively analyse data once new algorithms are available. Thus, we expect this type of methodological contribution to better support calls for reproducible science, and reduce data entropy through the use of open source software and raw data curation.

Machine vision is changing the nature and scope of the data collected by ecologists in two main areas. Firstly, digital sensors, digital raw data and the software used to analyse it have the advantage of providing replicable measurements with calculable accuracy and precision (see e.g. in Weinstein 2018), thus reducing observer bias. Moreover, digital raw data in conjunction with adequate meta-information and appropriate storage is less susceptible to entropy (i.e. decay or degradation Michener 2006; Hart *et al.* 2016) and falsification than data condensations (e.g. spreadsheets) while maintaining data dimensionality. Secondly, it can create opportunities for collaborations between ecologists and other scientific disciplines (e.g. engineers, computer scientists as proposed by Weinstein 2018) and with appropriate incentives will promote and advance data sharing and open-data practices, which will strengthen ecological research. Preserving raw data from instruments is paramount so data can be revisited and reanalysed under new paradigms or tools. But, we echo the sentiment of Veiga *et al.* (2017) around concerns of using data sets beyond their intended scope and breadth and data quality recommendations. For scientists relying on visual observations of organisms or specimens, optical sensors and current advances in machine vision are proving to be useful in opening new possibilities to move ecological research forward (e.g. Matai *et al.* 2012; Dell *et al.* 2014; Konovalov *et al.* 2019; Piechaud *et al.* 2019; Schneider *et al.* 2019). Machine vision and learning has the potential to revolutionize our current sampling methods and analysis in such a way that allow us to address rapidly and efficiently the urgency of sampling natural systems intensively and extensively, hence improving our understanding of these systems and our ability to manage them.

## Acknowledgements

We are thankful to K Sambrook and K Johns for editorial help on early versions of this manuscript. We want to acknowledge the Advanced Scientific Programming in Python Asia Pacific summer school for a travel grant and the training conferred to CH. CH is grateful to the Graduate Research School, James Cook University for their support during his PhD candidature. We thank A Barnett, K Abrantes, K Sambrook, A Hernandez, N Andrade and the Estuary and Coastal Wetland Ecology Research Group for invaluable discussions and suggestions in the making of this manuscript.

## Authors’ contributions

All authors conceived the experiment. CH performed fieldwork, programming and analysis. MS and RB contributed materials and funds. CH wrote the first draft, and all authors edited and reviewed the manuscript.

## Data accessibility

Crabspy – the toolbox used for extracting data and the R code used to analyse data and create figures are available in GitHub (https://github.com/CexyNature/Crabspy and https://github.com/CexyNature/CrabsMotionAndNetworks). Raw data used in this manuscript can be downloaded from the Tropical Data Hub (link to data would be published after acceptance in peer reviewed journal). Supplementary material can be found in Figshare (https://doi.org/10.6084/m9.figshare.12284435.v1).

## Funding

This research was supported by an Australian Government Research Training Program (RTP) Scholarship to CH. The study was funded by the Holsworth Wildlife Research Endowment - Equity Trustees Charitable Foundation & the Ecological Society of Australia, and College of Science and Engineering Joint research training grant, James Cook University.

## Notes

https://doi.org/10.5281/zenodo.3820270

https://doi.org/10.6084/m9.figshare.12284435.v1

https://github.com/CexyNature/CrabsMotionAndNetworks

### Competing Interest Statement

The authors have declared no competing interest.

